# Integrated metabolomics data analysis to generate mechanistic hypotheses with MetaProViz

**DOI:** 10.1101/2025.08.18.670781

**Authors:** Christina Schmidt, Denes Turei, Dimitrios Prymidis, Macabe Daley, Christian Frezza, Julio Saez-Rodriguez

## Abstract

With the growing number of metabolomics and lipidomics studies, robust strategies for bioinformatic analyses are increasingly important. However, the absence of standardized and reproducible workflows, coupled with ambiguous metabolite annotations, hampers effective analysis, particularly when integrating prior knowledge with metabolomics data. Moreover, the limited availability of comprehensive, curated prior knowledge further limits functional analyses and reduces the extraction of meaningful biological insights.

Here we present MetaProViz (Metabolomics Processing, functional analysis and Visualization), a free open-source R package for metabolomics data analysis that integrates prior knowledge to generate mechanistic hypotheses (https://saezlab.github.io/MetaProViz/). MetaProViz offers a flexible framework consisting of five modules: processing, differential analysis, prior knowledge integration, functional analysis and visualisation, applicable to both intracellular and exometabolomics experiments. To improve functional analysis, we created the Metabolism Signature Database (MetSigDB), a collection of annotated metabolite sets. MetSigDB includes classical pathway-metabolite sets, metabolite-receptor and metabolite-transporter sets, and chemical class-metabolite sets. In addition, MetaProViz enables the conversion of gene sets to metabolite sets by using enzyme-metabolic reaction associations. In addition, MetaProViz translates between metabolite identifiers of commonly used databases, analyzes mapping ambiguities and completes missing annotations. The MetaProViz functional analysis toolkit includes sample metadata analysis, classical enrichment analysis and biologically informed clustering. We showcase MetaProViz functionalities using kidney cancer metabolomics data from cell lines, cell-culture media, and tumour tissue. We found increased methionine usage in clear-cell renal cell carcinoma (ccRCC) cell lines in line with decreased methionine levels in tumour samples. Further, we link this observation to enzymes and transporters crucial for overall survival in ccRCC and suggest that the increased methionine usage reflects the elevated DNA-hypermethylation landscape, a known characteristic in ccRCC. In summary, MetaProViz facilitates and improves the analysis and interpretation of metabolomics data.

**Graphical Abstract:** 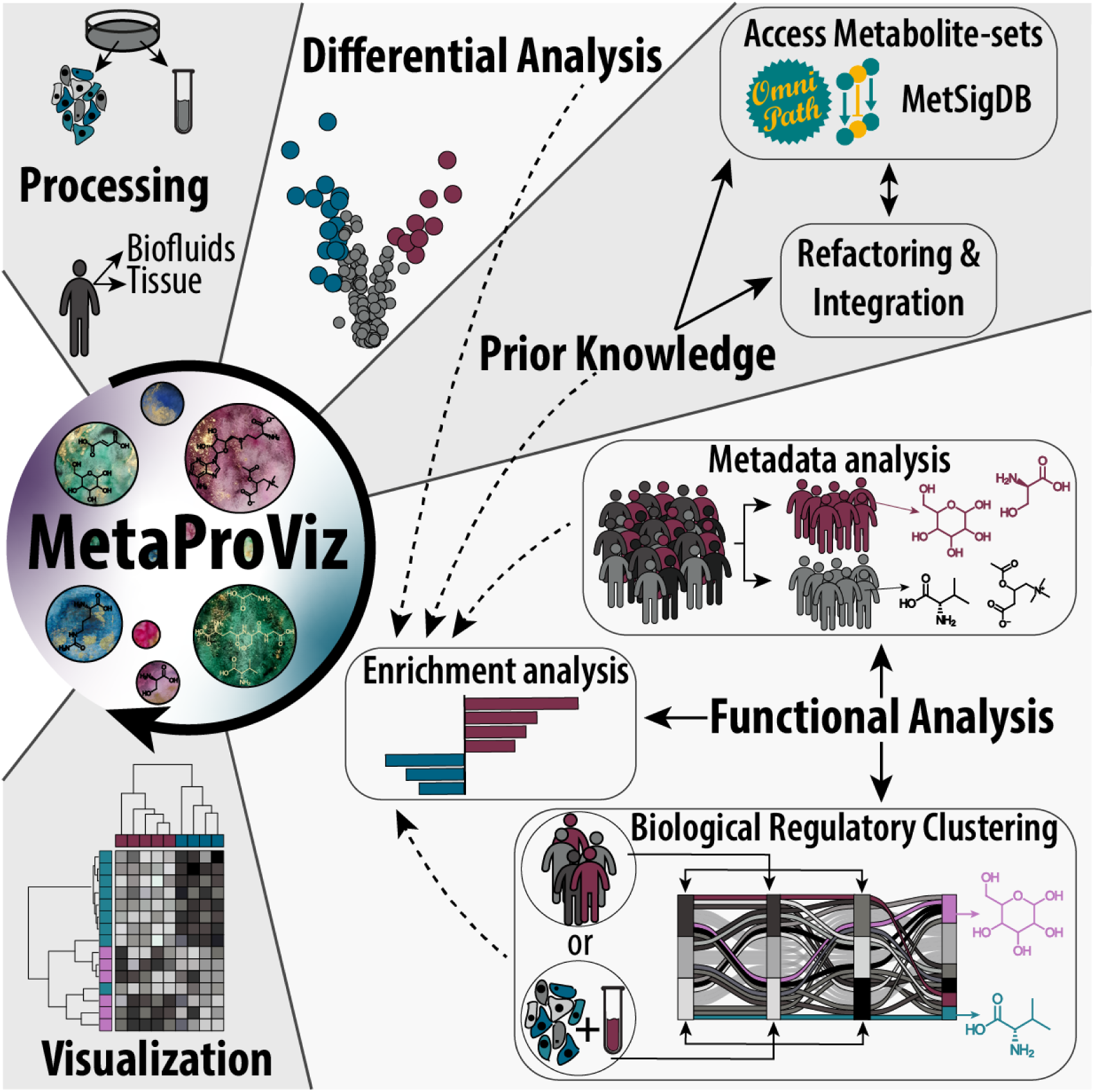

## INTRODUCTION

Advances in liquid-chromatography mass spectrometry (LC-MS) enabled the detection of thousands of metabolites from single biological samples, greatly expanding the depth of metabolomics and lipidomics datasets^1^.

A broad range of tools have been developed to facilitate different steps in metabolomics and lipidomics data analysis^2^. The analysis of metabolomics data encompasses several steps starting with pre-processing (including peak picking, feature assembly and -annotation, feature filtering, etc.), followed by normalisation of intensities and statistical analysis, culminating in functional and mechanistic analyses to ultimately interpret the data^3^. Examples of popular tools include CAMERA^4^, XCMS2^5^, and CliqueMS^6^ for the assembly, visualisation, and annotation of features. Some of them are also organised into workflows by other tools, for example MetaboAnalyst uses XCMS2 for peak-integration and CAMERA for peak-annotation^7,8^. Moreover, there are tools that focus on specific steps of the pre-processing of feature intensities, which encompasses feature selection, missing value imputation (MVI) and data normalisation. For example, MetImp4 is a web-tool that includes and compares multiple MVI methods^9^. The variety of MVI, normalisation and quality control techniques complicates the choice of methods that maximise data utility while minimising bias^2^. In experimental design, quality control samples are key for an effective removal of a large part of the technical variation^10^. Yet, many tools lack implementations or reasonable defaults for these critical steps, ultimately requiring advanced expertise to navigate the analysis landscape. Furthermore, many tools implement workflows^2^, typically spanning from feature annotation to functional analysis. However, most workflows are only accessible as web apps^2^, with the notable exception of MetaboAnalystR 4.0^11^. Their user-friendly interfaces often hide key settings, hindering compliance with today’s reproducibility standards^12^.

Functional interpretation is further hindered by the lack of unambiguous and reproducible metabolite annotations^13^. Since high-throughput MS methods are unable to distinguish fine structural details (e.g. stereoisomers or constitutional isomers)^14–16^, features do not correspond to fully defined structures. This is in discrepancy with assigning one particular, unambiguous IUPAC (International Union of Pure and Applied Chemistry) name or InChI (International Chemical Identifier) in addition to metabolite identifiers (IDs) as recommended by the Metabolomics Standards Initiative (MSI)^14–16^. Ultimately, this necessitates the assignment of multiple database entries to one feature, or one database entry representing all possible structures at once, since databases can have entries with different degrees of ambiguity^17^ (for example, CHEBI:16449 covers all varieties of alanine). This complexity in metabolite identifier assignment makes it difficult to link prior knowledge to features. Coupled with the challenge of defining the background set^18^, this undermines the robustness of enrichment analysis, like over representation analysis (ORA). Moreover, the distinct nature of metabolites compared to genes often render standard analytical approaches inappropriate, leading to misleading or nonsensical results^19^.

Here we present MetaProViz, **Meta**bolomics **Pr**ocessing, functi**o**nal analysis and **Vi**suali**z**ation, an R package to facilitate the analysis and biological interpretation of metabolomics data. MetaProViz covers processing (i.e. total ion count normalisation, feature filtering, etc.) and quality control, differential metabolite analysis, prior knowledge integration, functional analysis and visualisation. MetaProViz includes a collection of annotated metabolite sets named Metabolism Signature Database (MetSigDB) with classical pathway-metabolite sets, chemical class-metabolite sets, metabolite-receptor and metabolite-transporter sets. Furthermore, it enables the conversion of gene sets to metabolite sets. MetaProViz translates between metabolite IDs, defines mapping ambiguities between metabolite IDs of different databases and adds additional synonym IDs to identified features. MetaProViz also supports exometabolomics, the measurement of consumption and release of metabolites by cells^20,21^. Together with functional analysis, these tools help harness the potential of metabolomics data by enabling the generation of mechanistic hypotheses and improving biological interpretability. Through safeguards like automated checks and informative messages MetaProViz addresses common pitfalls, improves decision-making and prevents mis- and overinterpretation, ultimately making metabolomics analysis more accessible.

## RESULTS

### The MetaProViz software toolbox

MetaProViz consists of five main modules: 1) pre-processing, 2) differential metabolite analysis (DMA), 3) prior knowledge access, integration and optimization, 4) functional analysis, and 5) visualisation (**Fig. 1**). Each component can be used interactively or as building blocks within larger workflows. MetaProViz requires annotated and quantified features from peak integration as input, i.e. a matrix of intensities with annotated metabolites as column names and samples as row names. This format is commonly provided by facilities following feature assembly and annotation tailored to their extraction methods, instrumentation, analytes, and internal standards.

**Figure 1:**
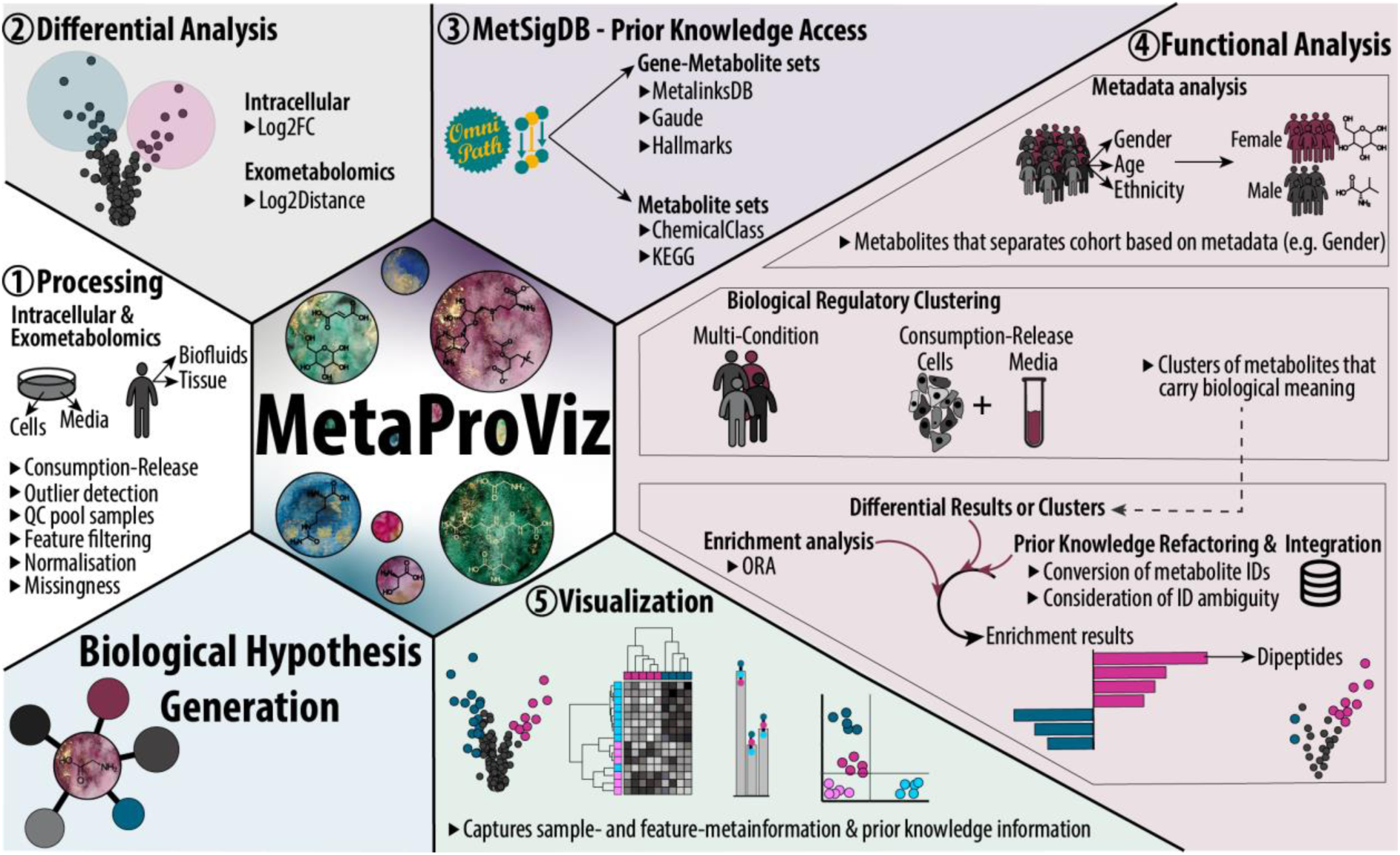
The contents and functionalities of MetaProViz. Overview of the five MetaProViz modules (1-5) and their main functionalities to ultimately aid biological hypothesis generation. Results of modules 1-4 can be visualized using the visualisation module 5. Differential results from module 2 and prior knowledge from module 3 can be used together within module four to perform over representation analysis. All modules can be used interactively as building blocks for a pipeline. Abbreviations: Principal Component Analysis (PCA), Quality Control (QC), Fold Change (FC), identifier (ID).

To guide the user, MetaProViz features different vignettes with example data, logs and return user messages, enabling even inexperienced users to make informed decisions. MetaProViz example datasets cover different experimental contexts, offering a robust foundation for testing and validating analysis workflows. The datasets have been sourced from kidney cancer studies (**SFig.1a**): (1) intracellular metabolomics of human renal epithelial cells (HK2) and clear-cell renal cell carcinoma (ccRCC) cell lines 786-O, 786-M1A, and 786-M2A, (2) corresponding cell culture media profiling (exometabolomics), which additionally includes OS-RC-2, OS-LM1, and RFX-631 ccRCC cell lines^22^ (**SFig. 1a**), (3) patient tissue metabolomics from 138 matched ccRCC and adjacent normal tissue pairs with extensive patient metadata^23^ (**SFig. 1b**), and (4) transcriptomics and proteomics data from ccRCC tissue^24,25^ (**SFig. 1c**) to showcase the integration of metabolomics with other omics data. During the analysis, MetaProViz ensures reproducibility by capturing parameters, operations, warnings, and errors in a log file, while reporting key parameters, best practices, and implications of analytical choices. All tables and figures are saved in the desired format, organized into subdirectories reflecting the workflow, and with easy-to-identify file names (**SFig. 1b**). These outputs include the final tables, intermediate results, prior knowledge and quality controls. The results are also returned as plot objects and data frames, enabling their downstream use. To create figures MetaProViz uses primarily the ggplot2 package^26^ with custom adjustments to labels, colours, and resolution for optimal slide and print quality. These publication-ready plots save users time by providing high-quality visualisations that meet professional standards.

MetaProViz has built-in access to prior knowledge adapted for enrichment analysis. This is done by systematically excluding non-specific metabolites (e.g. H_2_O or CO_2_), ions or xenobiotics. We refer to the collection of these metabolite sets as MetSigDB (**SFig. 1c**). This currently includes KEGG^27^ pathway-metabolite sets, chemical class-metabolite sets that classify chemical compounds into a hierarchical system based on their structural features^28^, MetalinksDB metabolite-receptor and metabolite-transporter sets^29^ and the ability to convert any gene-set into gene-metabolite-sets using metabolic reactions of a filtered version of Recon-3D accessed from the COSMOS prior knowledge network^30^. For the latter, we provide the Hallmarks pathway-gene sets^31^ and pathway-gene sets from Gaude et al.^32^ and convert those into pathway-gene-metabolite sets. The database knowledge in MetaProViz can also be used for other purposes, for example, the metabolite-receptor, metabolite-transporter, and enzyme-metabolite relationships for network analysis, and the ID translation utility to translate any data frame column with metabolite IDs.

### Improving the connection between prior knowledge and metabolomics features

Many databases that collect metabolite information, such as the Human Metabolome Database (HMDB)^33^, include multiple entries for the same metabolite with different degrees of ambiguity^17^. This poses a difficulty when assigning metabolite IDs to measured data where e.g. stereoisomers are not distinguished. Yet, if detection is unspecific, it is crucial to assign all possible IDs to increase the overlap with the prior knowledge. Since existing solutions such as MetaboAnalystR 4.0^11^ do not handle ambiguous metabolite id mapping, MetaProViz offers methodologies to solve such conflicts to allow robust mapping of experimental data with prior knowledge. Hence, MetaProViz assigns other potential metabolite IDs by translating IDs between databases using RaMP-DB^28^ in combination with a manually curated amino-acid collection. For features in the ccRCC patients samples, three different ID types have been provided by the authors^34^, namely HMDB, KEGG and PubChem. We found that most peptides only had a PubChem ID assigned (**Fig.2a**, left, yellow and **Supplementary Table 1**). By assigning other potential metabolite IDs and by translating between the present ID types, we not only increase the number of features within all ID types but also increase the feature space with HMDB and KEGG IDs (**Fig. 2a**, right, **SFig. 2 and Supplementary Table 1**). This is an important step to ensure mapping to metabolite-sets. Hence, when assigning an HMDB ID to alanine, assigning both IDs for D-alanine and L-alanine has to be done as no unspecific HMDB ID is available (**Fig. 2b, left** and **Supplementary Table 1**). On the contrary, when using ChEBI IDs, only one ID can be assigned, namely alanine (“CHEBI:16449”). However, this may lead to other problems as the ChEBI ID is not specific and does not map to most metabolic pathways in standard prior knowledge pathway-metabolite sets such as KEGG, WikiPathways and Reactome (**Fig. 2b, right** and **Supplementary Table 1**). Substrate chirality is critical to enzymatic processes and stereoselectivity of enzymes to be homochiral with predominance of one particular enantiomer (e.g. D-sugars, L-amino acids, etc.), hence it is important that a specific metabolite ID maps to certain pathways. An additional consideration when integrating measured metabolite IDs with prior knowledge is the inconsistency in identifier types across resources, for example, KEGG pathways use KEGG IDs, whereas Reactome relies on ChEBI identifiers. Hence, additional metabolite IDs often have to be assigned by translating metabolite IDs from one ID type into another (**SFig. 2** and **Supplementary Table 1**). Using RaMP-DB prior knowledge^28^ MetaProViz is able to translate between different ID types. During metabolite ID translation we can observe one-to-none, one-to-one, one-to-many and many-to-many mappings, which can happen either within a group, i.e. pathway, and/or across groups (**Fig. 2c-d** and **Supplementary Table 2**). Using MetaProViz’s functionality to summarize mapping ambiguities during ID translation, we demonstrate that converting KEGG IDs from KEGG pathway-metabolite sets to HMDB IDs results in zero one-to-one mappings. Instead, most mappings are one-to-many, and some even involve many-to-many relationships (**Fig. 2c**). This mapping ambiguity can lead to problems when set enrichment analysis is performed since it can inflate or deflate the pathway-metabolite sets. Therefore, MetaProViz maps the detected metabolites to the pathway-metabolite sets and flags ambiguous or problematic scenarios. This ensures that the number of features assigned to each pathway matches the original composition defined in the prior knowledge resource (**Fig. 2e**). In detail, there are scenarios that will require a reduction in features per pathway (**Fig. 2e**, Scenario 4) or the merge of two detected features by sum or mean (**Fig. 2e**, Scenario 5). Additionally, there are scenarios that only become apparent across pathways (**Fig. 2e**, Scenario 6-8) where decisions on a case-by-case basis are required.

**Figure 2:**
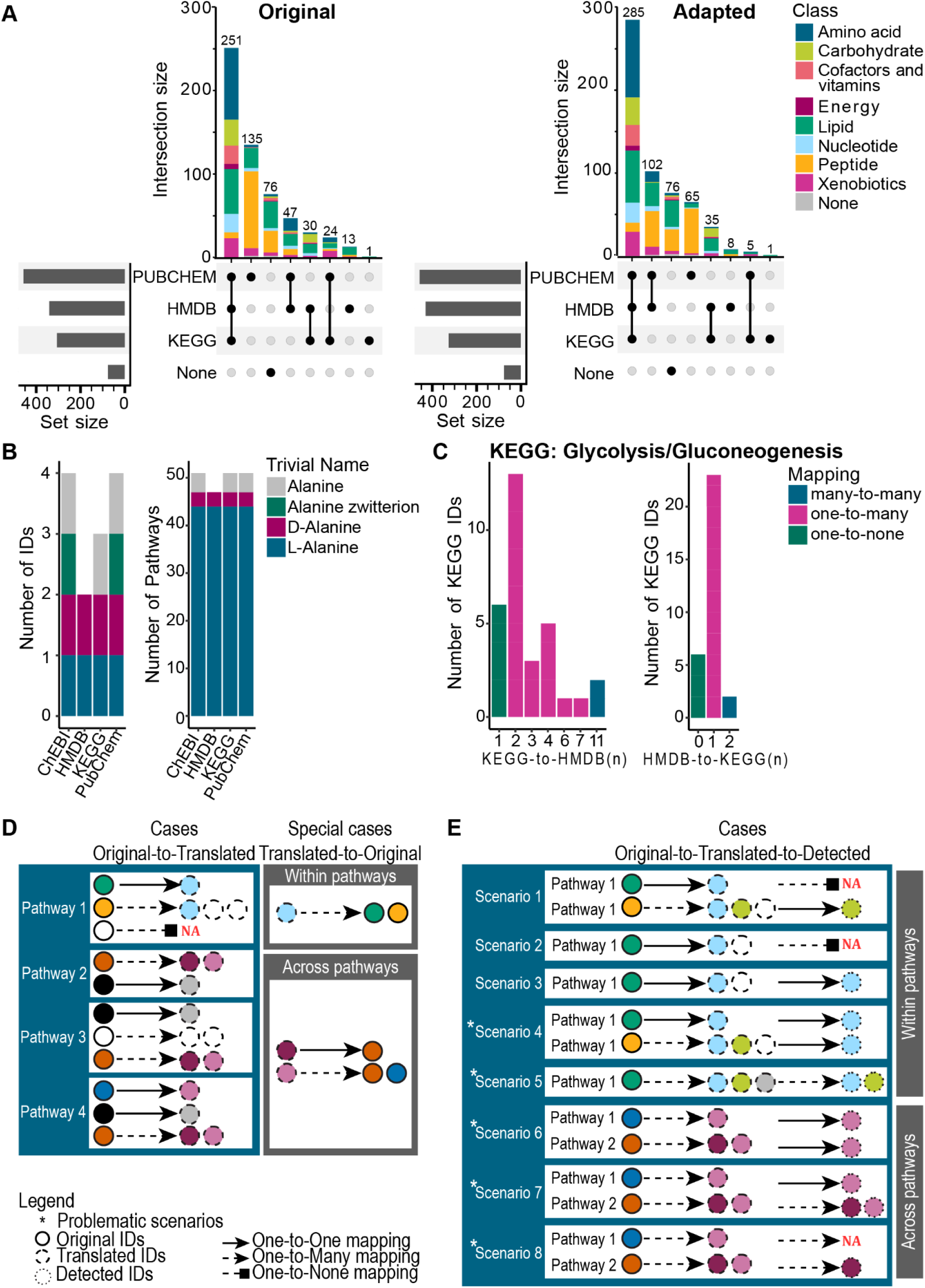
ID mapping problems in metabolite prior knowledge. **a)** Upset plots of the ccRCC patients feature identifiers as provided in the original publication (left) and after assigning additional potential IDs and translating between the provided ID types (right). **b)** Example of the presence of multiple metabolite IDs for the amino acid Alanine in ChEBI, HMDB, KEGG and PubChem databases (left) and the number of pathways these metabolite IDs can be mapped to by RaMP-DB (right). **c)** Ambiguity in the translation of metabolites in the “KEGG Glycolysis/Gluconeogenesis” pathway from KEGG to HMDB IDs (left) and then back (right) using MetaProViz; number of source IDs (bar heights) that map to certain number of target IDs (x-axis). **d-e)** Schematics of possible mapping cases between metabolite IDs (= each circle corresponds to one ID) of a pathway-metabolite set (e.g. KEGG) to metabolites IDs of a different database (e.g. HMDB) with **(d)** showing many-to-many mappings that can occur within and across pathway-metabolite sets and **(e)** additionally showing the mapping to metabolite IDs that were assigned to the detected peaks within and across pathway-metabolite sets.

In summary, the prior knowledge normalisation methods of MetaProViz increase the overlap between prior knowledge and measured data, leading to more accurate set enrichment results and improved functional interpretation. These functionalities include translating metabolite IDs, summarising mapping ambiguities that occur during ID translation and flagging problematic scenarios of mapping ambiguities (**Fig. 1**). Together with MetaProViz curated selection of annotated metabolite sets (**SFig. 1c**) and the implementation of ORA on differential analysis results or clusters of metabolites, these functionalities support the generation of mechanistic hypotheses.

### Preprocessing and quality control using MetaProViz

MetaProViz offers a suite of pre-processing and quality control (QC) functions, with a focus on user-guidance and reproducibility. This suite includes functions for both, the analysis of QC samples, such as pool samples, as well as functions for normalisation, missing value imputation and outlier detection. These steps collectively help remove low-quality features and sample outliers, resulting in high-quality data suitable for addressing biological questions. Here we apply MetaProViz functionalities to perform the pre-processing of exometabolomics data (**Fig. 3a**) from one renal control cell line and six ccRCC cell lines (**Fig. 3b**).

**Figure 3:**
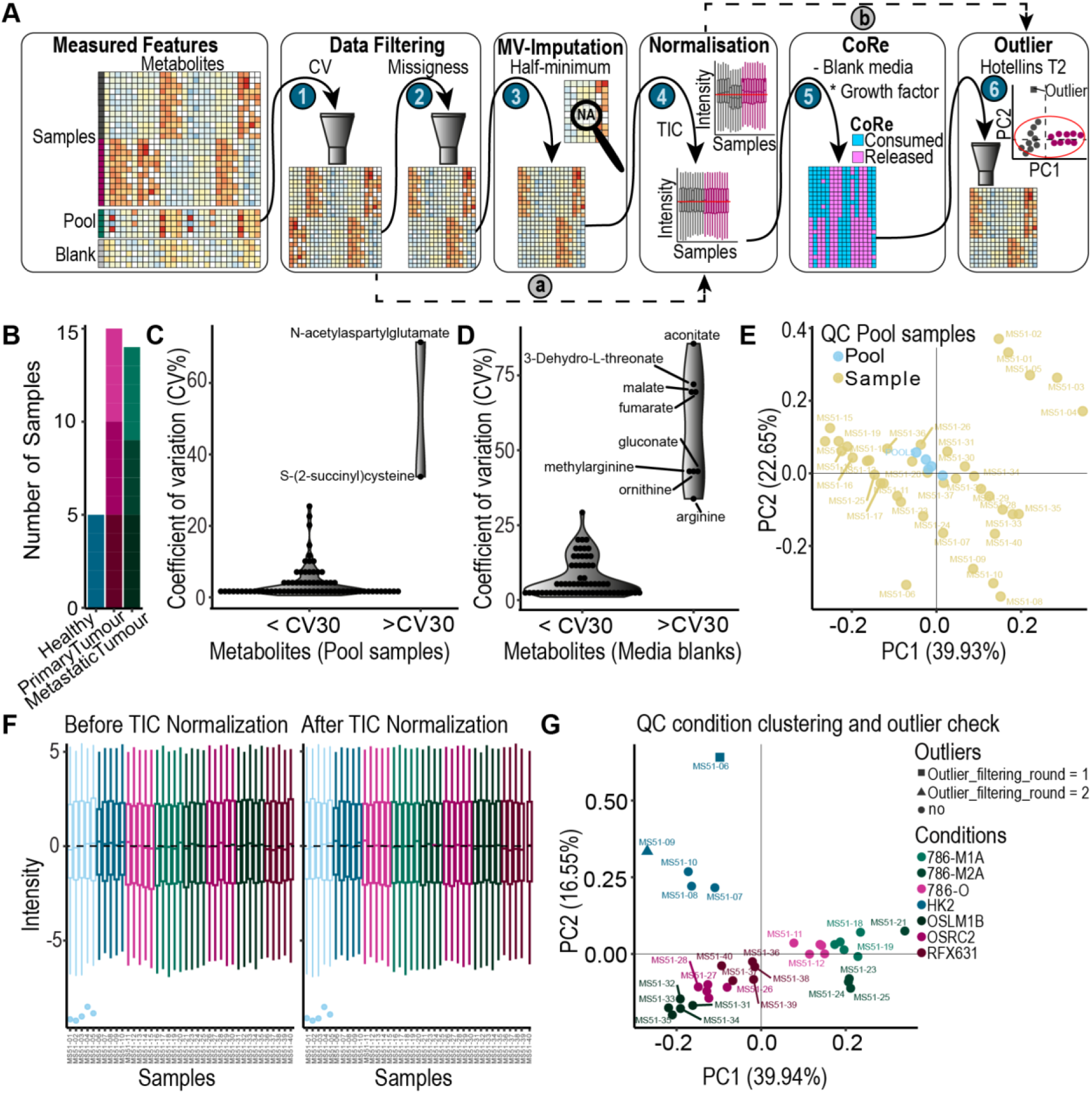
Quality control and processing of exometabolomics data using MetaProViz. **a)** Overview of the pre-processing workflow starting from a matrix of annotated features and divided into 6 steps: Features are filtered based on (1) coefficient of variation (CV) of pool samples and blanks and (2) missingness in the biological samples, (3) if applicable MVI using half-minimum is performed, (4) TIC normalisation, (5) if applicable Consumption-Release (CoRe) normalisation by subtraction of blank media samples and by multiplying with growth factor and (6) outlier removal based on Hotellin’s T2 distribution. It is common that MVI and CoRe are skipped (route (a) and (b)). **b)** Number of exometabolomics samples from ccRCC cell lines including healthy (blue: HK2), primary tumour (light to darkpink: 786-O, OSRC2, RFX631) and metastatic (light to darkgreen: 786-M1A, 786-M2A, OSLM1B). **c-e)** Exometabolomics feature quality assessment of: **c)** CV of each metabolite in pool samples, **d)** in media blank samples, which are samples where no cells were cultured in, and **e)** PCA showing metabolite pool samples clustering around the origin. **f)** Boxplots of feature intensities before and after TIC normalisation. **g)** PCA plot of Hotelling’s T2 outlier detection performed on the processed and normalised exometabolomics data using 99% confidence interval. Detailed explanation, code and plots can be found in the vignette (https://saezlab.github.io/MetaProViz/articles/CoRe%20Metabolomics.html).

Solutions to identify low-quality features based on QC samples are multifaceted. Quality control (QC) samples such as blanks (e.g., water or PBS), media blanks (media without cells), and pool samples (a homogeneous mixture of all samples), can be used to assess feature variability. Specifically, the coefficient of variation (CV) helps identify features with high variability and thus low detection accuracy. For blanks, minimal or no signal is expected, while media blanks and pool samples typically show CVs below 30% (**Fig. 3c-d, SFig. 3a-b** and **Supplementary Table 3**). In our data, most measured features have CVs near zero, indicating high measurement confidence (left violin **Fig. 3c-d**). Additionally, pool samples are expected to cluster near the coordinate origin in PCA plots, reflecting their role as a homogeneous mixture of all samples (**Fig. 3e**). Beyond removing low-quality features based on variability, features can also be filtered based on missingness, commonly following the 80% filtering rule in metabolomics research^9,35^. Here the missingness for each feature can be quantified across all samples^35^ or across samples per-condition basis, e.g. for each cell line^9^. From a biological standpoint, filtering out low-quality or highly missing features is essential to avoid skewing downstream analyses by inflating the importance of certain pathways dominated by unreliable measurements.

To prevent biases in metabolomics data caused by variations in sample loading or instrument performance, a common approach is to normalize each sample by its Total Ion Count (TIC)^36^. This involves dividing each feature intensity by the sample’s TIC and scaling it by the mean ion count across all samples (**Fig. 3f** and **Supplementary Table 3**). However, users are cautioned against applying TIC normalisation to datasets with a small number of features, as this may lead to inaccuracies^36^. Exometabolomics experiments require additional normalisation to allow interpretation of metabolite consumption or release^37^. Specifically, each feature is normalized using both media blanks and either a growth factor (to account for overall cell growth during the experiment) or growth rate (to capture variations in cell proliferation). This normalization yields negative values when a metabolite is consumed from the media, and positive values when it is released by the cells into the culture medium. This is crucial to understand metabolite exchange with the extracellular environment and can be used to study metabolic pathway activity^37^. Even though commonly used, Missing Value Imputation (MVI) can impact and potentially skew biological interpretation. Hence, we opted to only provide MVI for values that are missing not at random (MNAR), where half minimum MVI has been shown to perform best for metabolomics^9^. By performing PCA, we can check for batch effects, which are technical variations that confound downstream analysis, by e.g. colour coding for biological replicates (**SFig. 3c**). Given the multitude of factors that can introduce batch effects, and the challenges they pose when conflated with true biological signal^38^, we decided to not include batch correction algorithms in MetaProViz and encourage the user to apply dedicated tools for this. Once the data processing and normalisation is complete, Hotelling’s T2 test^39^ is used to detect outliers across the multidimensional space of PCA (**Fig. 3g, SFig. 3d-e**). It is up to the user to remove any sample outliers, but we recommend removing major outliers identified in the filtering round 1 (**Fig. 3g**). Here it can also be helpful to use biological information such as the experimental conditions of the cell lines in this example (**Fig. 3g and Supplementary Table 3**). In this case, it becomes clear that sample MS51-09, identified as potential outlier in filtering round 2, clusters together with replicates of the same cell line, hence we would opt to not remove it.

### Combination of exo-with intracellular metabolomics and prior knowledge to generate mechanistic hypothesis

To harness the full potential of exometabolomics data and unravel metabolic dysregulation, MetaProViz offers a suite of functional analysis tools that use differential metabolite analysis results as input. Starting from the normalized data matrix (normalisation details see methods and **Fig. 3**), we performed differential metabolite analysis using MetaProViz ccRCC cell line example data (**Fig. 3a**). Since exometabolomics data normalized to culture media blanks includes both positive and negative values, reflecting metabolite release and consumption, respectively, classical log2 fold change cannot be applied. Instead, we calculate the log2 distance (**Fig. 4a**, methods section), which captures whether a metabolite is released or consumed in both conditions, or released in one and consumed in the other (and vice versa). This information is visualized using a color-coded scheme (**Fig. 4a** and **Supplementary Table 3**). For statistical testing we used limma^40^ and one-versus-all comparisons with p-value adjustment using false-discovery-rate (fdr). To interpret exometabolomics data of multiple conditions, we generate feature metadata that summarises the consumption-release information based on the mean of the measured replicates per condition. These summaries are visualized in a heatmap, which displays cell line clustering according to their metabolite consumption-release profiles (**Fig. 4b**). To aid interpretation, we selected a representative subset of cell lines, healthy (HK2), primary ccRCC (786-O), and metastatic ccRCC (786-M1A), and selected metabolites based on their assigned primary pathway. Focusing on amino acid metabolism, we found that primary and metastatic ccRCC cell lines (786-O and 786-M1A, respectively) predominantly consume various amino acids whilst healthy cells (HK2) produce and release those to the media (**Fig. 4c**).

**Figure 4:**
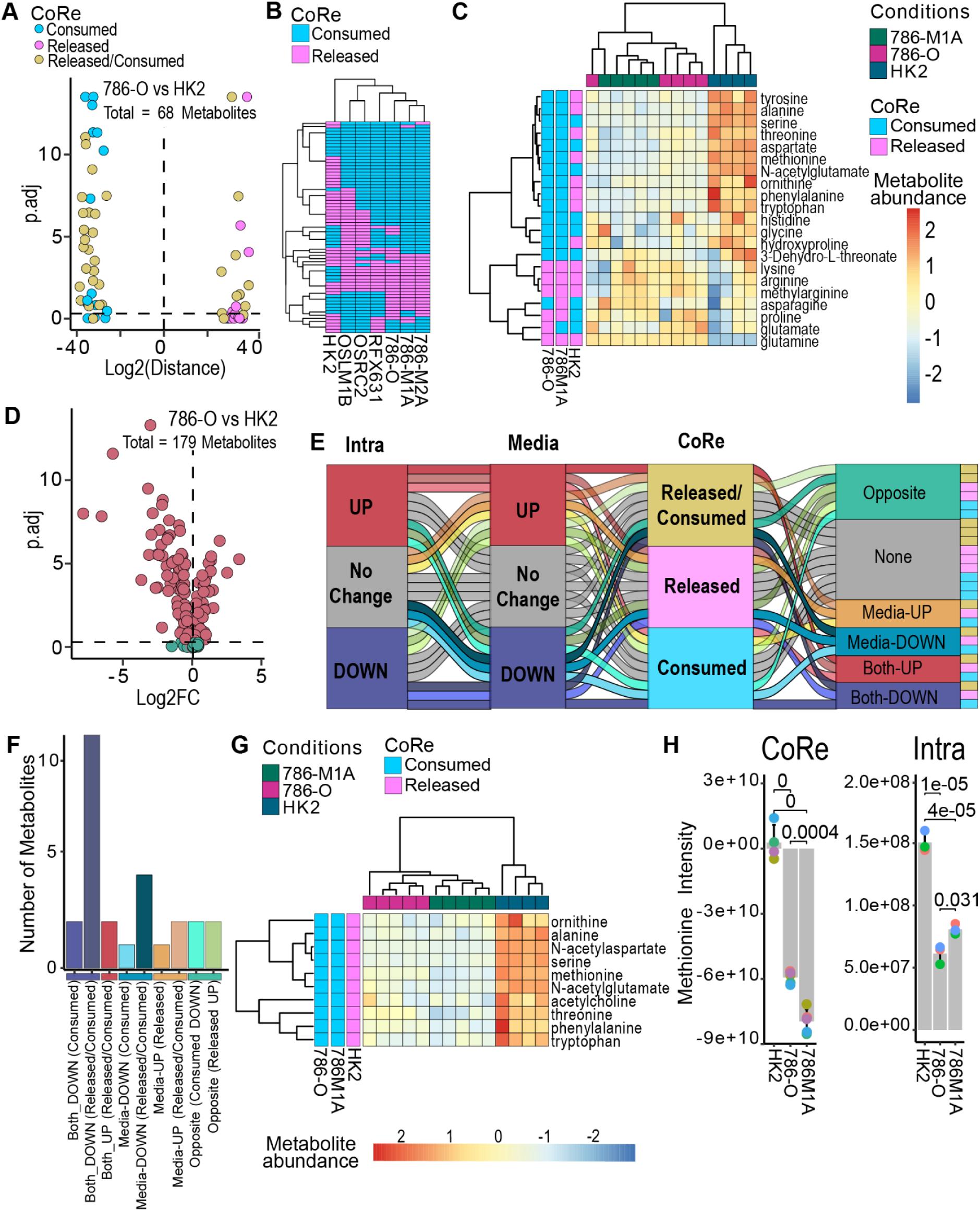
Combination of exo-with intracellular metabolomics and prior knowledge. **a)** Differential metabolite analysis results for exometabolomics data comparing 786-O versus HK2 cells. **b)** Heatmap of mean consumption-release of the measured metabolites across cell lines. **c)** Heatmap of normalised ccRCC cell line exometabolomics data for the selected metabolites of amino acid metabolism for a sample subset. **d)** Differential metabolite analysis results for intracellular data comparing 786-O versus HK2 cells. **e)** Schematics of bioRCM process to integrate exometabolomics with intracellular metabolomics and **f)** number of metabolites by their combined change patterns in intracellular- and exometabolomics in 786-M1A versus HK2. **g)** Heatmap of the metabolite abundances in the “Both_DOWN (Released/Comsumed)” cluster. **h)** Bar graphs of methionine intensity for exometabolomics (CoRe) and intracellular metabolomics (Intra).

Alongside exometabolomics experiments, intracellular metabolomics of the same cells is often performed. In MetaProViz we provide intracellular metabolomics for four of the seven cell lines (**Fig. 3a**), namely HK2, 786-O, 786-M1A and 786-M2A. To integrate exometabolomics with intracellular metabolomics, differential metabolite analysis results from the same comparison are required (**Fig. 4a, d** and **Supplementary Table 4**). These results serve as input for our biological regulatory clustering method (bioRCM), which is based on logical regulatory rules. We have previously demonstrated that bioRCM can extract meaningful feature clusters in a multi-omics context^25,41^. In MetaProViz, we created logical regulatory rules to integrate exometabolomics with intracellular metabolomics consisting of an ordered series of three 5-state-5-state-4-state transitions. The first layer includes the intracellular metabolomics, the second layer the exometabolomics and the third layer the direction (consumed, released, consumed/released or not detected) of the exometabolites (**Fig. 4e** and **Supplementary Table 5**). This will result in clusters of metabolites that carry biological meaning (**Fig. 4f** and **Supplementary Table 5**). Next, we focused on the cluster “Both_DOWN (Released-Consumed)” and found that several amino acids are consumed by the ccRCC cell line 786-M1A but released by healthy HK2 cells. At the same time, intracellular levels are significantly lower than in HK2 (Log2FC = −0.9, p.adj = 4.4e-5) (**Fig. 4g**). To explore the role of these metabolites in signaling, we queried MetalinksDB for their known upstream and downstream protein interactors for the measured metabolites (**Supplementary Table 5**). This approach is especially valuable for exometabolomics, as it allows us to generate hypotheses about cell– cell communication. Notably, we identified links involving methionine (**Fig. 4h**), enzymes such as *BHMT,* and transporters such as *SLC43A2* that were previously shown to be important in ccRCC^25,42^ (**Supplementary Table 5**). Interestingly, intracellular methionine levels comparing tumour versus normal tissue are decreased (**Fig. 4h**, right), and in the extracellular data methionine was consumed from the media whilst control cells released methionine (**Fig. 4h**, left). This increased methionine usage in ccRCC could be explained by the increased DNA-hypermethylation landscape, a known characteristic in ccRCC^24^.

In summary, calculating consumption-release and combining it with intracellular metabolomics and prior knowledge using the MetaproViz toolkit facilitates biological interpretation of the data.

### Understand metabolic alterations by patient metadata analysis and multi-condition clustering

Compared to cell line experiments, patient cohorts are generally more heterogeneous and come with extensive sample metadata (e.g. age, gender, therapeutic response or stage amongst many other information), which is also included with the ccRCC patient data in MetaProViz (**Fig. 5a**). To identify the main metabolite drivers, we created a method in MetaProViz that relates metabolite features to patient metadata. First this method orders patients by a PCA on their metabolomics features, then it performs multiple anova’s to quantify how much each principal component can be explained by each metadata variable. Applying this to the patient ccRCC data, as expected, tissue type (tumour or normal) was the strongest explanatory variable for PC1 (**Fig. 5b, Supplementary Table 1**). Another variable explaining the same amount of variance in PC1 is the tumour stage, which could point to adjacent normal tissue metabolic rewiring that happens in relation to stage. Interestingly, the variance in PC4 can be explained in 3.8 % by metabolites separating age and stage. We decided to dissect this further by applying bioRCM using the multi-condition clustering focusing on patient subsets of young (age <42 years) and old (age > 58 years) patients (**Fig. 5c**). The bioRCM we developed to interpret multi-condition experiments uses an ordered series of two 5-state-5-state transitions between the layers. The core states include “UP”, “DOWN” and “No Change” with the latter further subdivided into “Not detected”, “No significant change”, “Significant positive” and “Significant negative” (**Supplementary Table 4**). One of the most altered metabolite classes are dipeptides that could play a role as biomarkers in cancer^43^ and are upregulated in old patients, whilst downregulated or unchanged in young patients (**Fig. 5d**). Enrichment analysis using KEGG pathways further revealed that “Mineral absorption” and “Protein digestion and absorption” are specifically downregulated in young patients (**Fig. 4e, Supplementary Table 1**). In turn, this potentially explains the reduced number of dipeptides observed.

**Figure 5:**
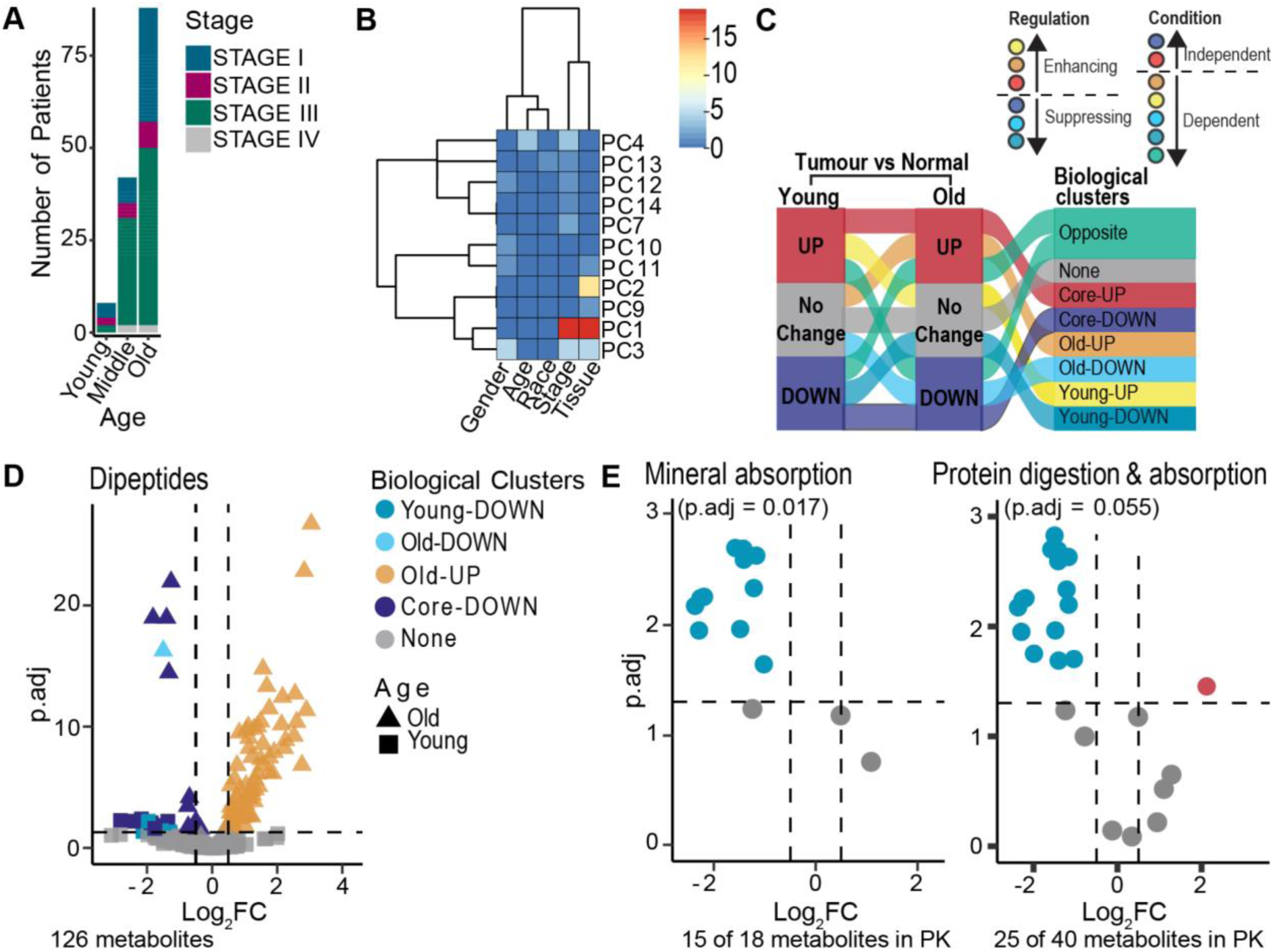
Patient metadata analysis and multi-condition clustering to dissect patient data. **a)** Number of patients in the ccRCC cohort by age at surgery <42 (young), >42 & <58 (middle) and >58 (old) and tumor stage. **b)** Heatmap of variance in principal components (PCs, y-axis) explained significantly by patient metadata (x-axis) (Tukey’s p.adj < 0.05). **c)** Schematics of two condition bioRCM process to compare changes in tumour versus normal in old versus young patients. It results in biological clusters that carry information about regulation and condition dependence (colours). **d)** Volcano plot of tumour versus normal metabolite changes in old (triangle) and young (square) ccRCC patients colour coded by biological clusters. **e)** Volcano plots of tumour versus normal metabolite changes in young patients within two KEGG pathways that were significantly deregulated based on over representation analysis.

As this example shows, performing metadata driven analysis can extract metabolic patterns of patient subgroups, which classical pathway enrichment analysis would not have been able to extract.

## DISCUSSION

MetaProViz is a modular, flexible, open-source and reproducible metabolomics analysis package that integrates curated prior knowledge with a rich functional analysis toolkit. Its capabilities extend beyond standard workflows by addressing key limitations in the field arising from the ambiguity of metabolite features and mapping of metabolite identifiers across databases. MetaProViz achieves this through dedicated tools that complete feature annotations with missing identifiers and quantifies mapping ambiguities, facilitating a more accurate and comprehensive integration of metabolomics data with existing knowledge. Ultimately this improves pathway enrichment and functional analyses, surpassing the identifier handling and enrichment options available in other metabolomics analysis solutions^11^. Besides intracellular metabolomics, MetaProViz also features dedicated methods for exometabolomics analysis, which, to our knowledge, no other publicly available tool currently provides. These methods enable the systematic analysis of metabolite consumption and release by cells and linking extracellular metabolic changes to intracellular pathways and mechanisms, expanding the scope of functional and mechanistic interpretation. Combined with visualisation features designed to minimise the risk of misinterpretation, MetaProViz supports the generation of mechanistic hypotheses and strengthens the biological interpretability of metabolomics studies.

The integration of metabolomics data with prior knowledge remains challenging due to ambiguity in feature identification and ambiguous mapping between ID types. There is no single solution that fits all problems, therefore, MetaProViz offers a set of functions to expand the metabolite ID space within datasets, translate IDs of data or metabolite-sets, quantify mapping ambiguities, and highlight problematic cases. We anticipate that future advances will move beyond strict reliance on database specific identifiers and leverage chemical structures and ontologies to explicitly resolve ambiguities, thus further improving mapping accuracy and interpretability. MetaProViz is already structured to accommodate such developments by using the OmniPath ecosystem^44^ at the backend, which enables us to address these limitations inherent to current ID-based methods in the future.

MetaProViz inclusion of prior knowledge of receptors, transporters and other protein interactors, provided by MetalinksDB, extends analysis beyond classical pathways by linking altered metabolite levels to signalling cascades. Applying this to clear-cell renal cell carcinoma (ccRCC) data, we identified increased methionine usage in ccRCC cell lines, and through MetalinksDB, linked this change to enzymes and transporters associated with overall survival^25,42^. Consistent with these findings, patient tumour samples exhibited reduced methionine levels compared to normal tissue, potentially reflecting the elevated DNA-hypermethylation landscape characteristic of ccRCC^24^. This case study illustrates the potential of MetaProViz’ to integrate metabolomics data with prior knowledge, to generate biologically meaningful hypotheses, and to place metabolic changes into a broader mechanistic context.

Metabolomics poses unique analytical challenges compared to genomics, and we designed MetaProViz to actively guide users in selecting appropriate methods. Significant development effort has gone into implementing informative messages and automated checks, for example, performing Shapiro–Wilk tests during differential analysis to report whether the chosen statistical test is appropriate. These safeguards help prevent common analytical pitfalls and support informed decision-making for both experienced bioinformaticians and researchers less familiar with metabolomics-specific issues. At the same time, the modular architecture of MetaProViz supports integration with complementary tools, such as COSMOS^30^ for multi-omics network analysis, and facilitates future extensions for single-cell and spatial metabolomics. Incorporation into Bioconductor^45^ and workflow management systems, such as nf-core^46^ modules in Nextflow^47^, will enhance the support of reproducible, scalable applications in metabolomics core facilities. Beyond current capabilities, future developments could also incorporate mechanistic modeling to capture metabolic fluxes, subcellular compartmentalization, enzyme kinetics, regulatory feedback loops, and thermodynamic constraints to dissect metabolic response under perturbations.

In summary, MetaProViz combines prior knowledge access, advanced identifier translation, and mapping ambiguity quantification with, exometabolomics analysis, functional analysis, and visualisation within a flexible and interoperable framework. Its unique support for exometabolomics, together with safeguards such as automated checks and informative messages address common pitfalls and improves decision-making during analysis. Its design balances the needs of bioinformaticians accustomed to genomic workflows but new to metabolomics, while also enabling wet-lab scientists with minimal coding experience to perform robust and reproducible analyses.

## FEATURES AND IMPLEMENTATION

### Example Data

MetaProViz provides access to diverse example datasets from kidney cancer studies. Cell-line data include intracellular metabolomics of human renal epithelial cells (HK2) and renal cell carcinoma (ccRCC) cell lines 786-O, 786-M1A, and 786-M2A and corresponding cell culture media profiling (exometabolomics), which additionally includes ccRCC cell lines OS-RC-2, OS-LM1, and RFX-631 were downloaded from metabolomics workbench project PR001418 including study ST002224 and ST002226^22^. Patient tissue metabolomics from 138 matched ccRCC and adjacent normal tissue pairs were downloaded from Hakimi et al. “supplementary table 2”, which includes median normalized data^34^. Patient tissue processed transcriptomics and proteomics data from ccRCC tissue were downloaded from Mora&Schmidt et al table “13073_2024_1415_MOESM3_ESM.xlsx” originally from proteomics Data Commons accession number PDC000127^24,25^. The detailed code can be found on the “ExampleData” branch of the MetaProViz package (https://github.com/saezlab/MetaProViz/tree/ExampleData).

### Data cleaning

To exclude measured features of low quality, the missingness for each feature across all samples per-condition was quantified (modified 80%-filtering rule)^9^ and features with missingness <= 20% were kept. Additionally features with CV >30% in the pool samples (homogenous mixture of all samples) were removed. Lastly, for exometabolomics features with CV>30 in media blanks (media without cells) were flanked.

### Data Processing

Total Ion Count (TIC) normalisation according to Wulff et al.^36^ was performed by dividing feature intensities by the sample’s total ion count and scaling by the mean ion count across all samples and thus ensures that variations in sample loading or instrumentation do not bias the data. For the exometabolomics experiments media sample (cell culture media taken from cells) were additionally normalised using both media blanks (samples where no cells were cultured in) and growth factor (accounts for cell growth during experiment) as growth rate (accounts for variations in cell proliferation) has not been available.

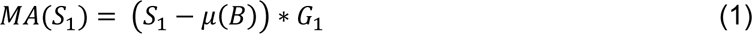

- MA represents the metabolite abundance (arbitrary unit) of a measured metabolite.
- *S_1_* represents sample 1.
- μ(*B)* represents the population mean of the blank samples.
- G_1_ represents the growth factor or growth rate corresponding to S_1_.

The results are either a negative value, if the metabolite has been consumed from the media, or a positive value, if the metabolite has been released from the cell into the culture media.

### Missing Value Imputation

For cell-line data we performed half minimum (HM) missing value imputation per feature assuming missing values that are missing not at random (MNAR) for which HM has been shown to perform best (Wei et al. 2018). Noteworthy, MetaProViz does not offer other ways to perform missing value imputation and urges the user to use dedicated tools for this.

### Outlier detection

Testing for outliers was performed using the Hotelling’s T2 outlier test (Hotelling 1931) with a confidence interval of 0.99. If an outlier was identified, it was removed, and the test was repeated. All results can be found in the supplementary tables.

### Differential Metabolite Analysis (DMA)

For statistical testing pairwise comparisons of conditions were defined by the metadata annotations and passed using numerator and denominator. For cell lines we performed one-vs-all comparisons (HK2 cells versus all other cells) using Anova. In general, MetaProViz supports all-vs-all one-vs-all and one-vs-one comparisons. For the latter, methods including Student’s t-test, Wilcoxon rank-sum test^48^, chi-squared test, or correlation analysis can be used. For one-vs-all or all-vs-all comparisons, methods such as ANOVA, Welch’s ANOVA^49^, Kruskal-Wallis test^50^, or linear modelling (via limma’s `lmFit` method^40^) are available. To account for multiple hypothesis testing, p-values were adjusted using the false discovery rate (FDR)^51^ method unless otherwise specified. Yet, MetaProViz also offers other adjustment methods such as Benjamini Hochberg (BH)^52^. Assessment of statistical assumption was done for each metabolite by checking the normality of metabolite distributions across conditions using the Shapiro-Wilk test^50^ and by evaluating the homogeneity of variances using Bartlett’s test^53^. For “**Standard DMA”** applied to intracellular metabolomics, the Log2 Fold Change (Log2FC) comparing Condition 1 (C_1_) versus Condition 2 (C_2_) is calculated based on the mean (average) of the sample values corresponding to the same condition C_n_:

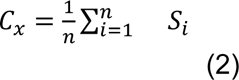

- C_x_ represents the condition a set of samples (S_i_) corresponds to.
- 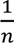 represents the reciprocal of the total number of values (n), which calculates the mean.
- 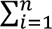 *S_i_* represents the summation of all the sample values, from S_1_ to S_n_.

Log2 Fold Change (Log2FC) comparing Condition 1 (C1) versus Condition 2 (C2)

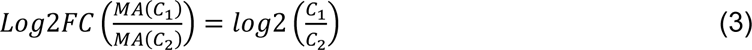

- 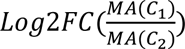 represents the Log2FC comparing the MA of C_1_ versus the MA of C_2_.

Statistical results of hypothesis testing for the difference between the individual sample MA values of C_1_ and the individual sample MA values of C_2_ is calculated using standard statistical tests as described above. For **“CoRe DMA”**, which is applied to exometabolomics, the absolute difference between C_1_ versus C_2_ was calculated by first calculating the mean (average) of the sample values corresponding to the same condition C_n_ using (1). In the next step we used C_x_ to calculate the absolute difference. In this step the absolute difference is log_2_ transformed to make the values comparable between the different metabolites.

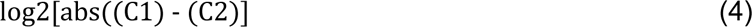

The result doesn’t consider whether one product is larger than the other; it only looks at the magnitude of their difference. To reflect the direction of change between the two conditions we multiplied with −1 if C1 < C2.

### Prior Knowledge access

MetaProViz uses the OmniPath database and the R package OmnipathR to access prior knowledge^44^. To support the applications in MetaProViz, we implemented new clients in OmnipathR for HMDB^33,54^, RaMP DB^28^, KEGG^27,55^, Hallmarks^31^, MetalinksDB^29^ and Gaude^32^. These clients import data directly from the original source in bulk and parse it into data frames. OmnipathR provides infrastructure for the clients, such as download management and local cache.

### Metabolite ID translation and addition

In MetaProViz, we make use of OmnipathR’s^44^ identifier translation utility, which is able to translate between all compound ID types supported by RaMP^28^ or HMDB^33,54^, in addition to several protein, gene and RNA ID types. We use this translation service also for adding additional potential metabolite IDs in combination with a manually curated amino acid table. The translation is highly customizable, and optionally an ambiguity analysis is carried out, qualifying and quantifying the one-to-one, one-to-many and many-to-many mappings in the data frame, which is summarised in the summary report. If grouping variables such as pathway names in pathway-metabolite sets are present, also ambiguities within and across groups were analysed and quantified.

### Prior Knowledge Refactoring and Contextualisation

To combine the exometabolomics data with MetalinksDB^29^, we contextualised MetalinksDB to only contain cell_location = “Extracellular”, tissue_location = c(“Kidney”, “All Tissues”) and biospecimen_location = c(“Blood”, “Urine”). Furthermore, we excluded all metabolites that were not part of the bioRCM cluster RG2_Significant “Both_DOWN (Released/Consumed)” (see biological regulatory clustering method for details on clustering).

### Biological Regulatory Clustering Model (bioRCM)

This is conceptually based on the Signature Regulatory Clustering (SiRCle)^25^ and the input data are differential metabolite analysis results including statistics and fold change or distance and direction for exometabolomics data. Depending on the input data available there are two bioRCM methods available, Multi-Condition bioRCM and Consumption-Release bioRCM. For Multi-Condition bioRCM multiple experimental conditions or patient metainformation have to be available that can be used to create multiple comparisons. Here we performed two differential analyses comparing tumour versus normal first for a subset of old patients (age at surgery >58) and second for a subset of young patients (age at surgery <42) of the patient’s example data, which was used as input for the bioRCM Multi-Condition clustering. For Consumption-Release bioRCM intracellular and exometabolomics of the same conditions are required. Here we performed differential analysis comparing 786-M1A versus HK2 cells using the intracellular and exometabolomics cell line example data.

The user will need to choose the “background method (BG)”, the thresholds/cutoffs of the input data and the “Regulation Grouping (RG)”. The BG setting defines which features are considered for the bioRCM clusters (**Table 1**). For example, Intra&CoRe is the most restrictive setting as only features detected in exometabolomics and intracellular metabolomics are considered. Here we have used the BG settings “C1&C2” for the multi-condition approach and “Intra&CoRe” for the Consumption-Release approach.

**Table 1:**
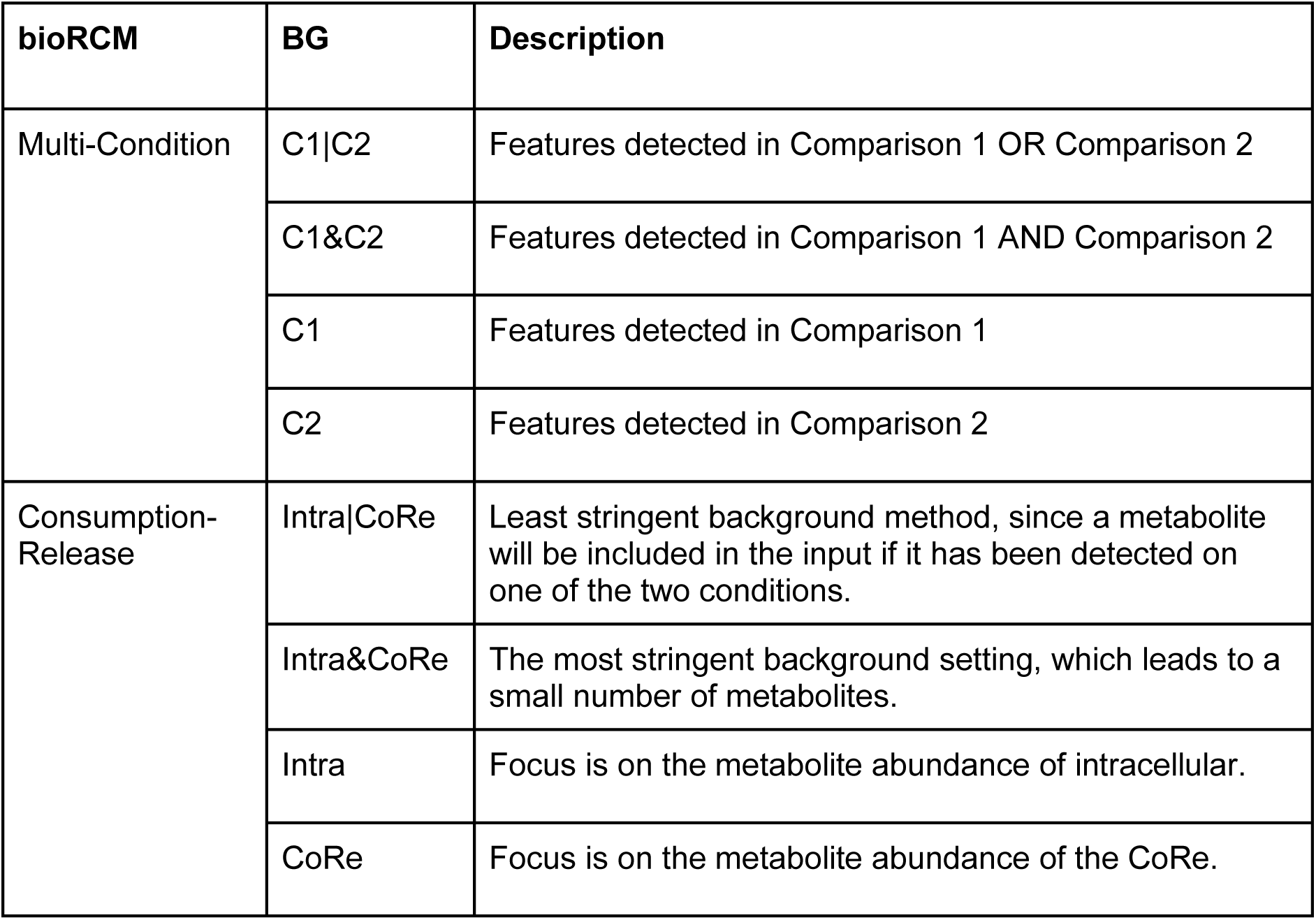
Overview of bioRCM background (BG) options.

Next, the user will set the different input thresholds, which will define whether a feature is considered “UP”, “DOWN” or “No Change”, with “UP”/ “DOWN” meaning a feature is significantly up-/downregulated in the underlying comparison and “No Change” are features that do not meet the significance and/or the fold change/distance threshold, or dependent on the background (BG) setting were not detected. Hence, “No Change” is subdivided into “undetected”, which includes features detected in one comparison/one data layer, but not in the other comparison/data layer, “not significant”, which includes features that do not meet the significance threshold set, “significant negative”, and “significant positive”, which includes features that do not meet the negative/positive fold change threshold but meet the significance threshold.

Lastly, the “Regulation Grouping (RG)” defines how the different flows will be summarised into clusters, whereby all flows can be found in RG1_All. The Multi-Condition bioRCM uses an ordered series of two 5-state-5-state transitions between the two layers (=comparisons), where the five states divide the metabolites into up-regulated, down-regulated, “significant negative”, “significant positive”, “undetected”. Similarly, also the Consumption-Release bioRCM includes these 5-state-5-state transitions for intracellular and exometabolomics, but additionally extends to a 5-state-5-state-4-state between the layers where the first layer includes the intracellular metabolomics, the second layer the exometabolomics and the third layer the direction (consumed, released, consumed/released or not detected) of the exometabolites. Those flows are summarised either with a focus on significant changes, meaning even metabolites that are UP, and significant positive will be placed in the same cluster, or with a focus on the change, meaning in this case UP and significant positive will not be placed in the same cluster. For the exact flows also check **Supplementary Table 4 and 5**.

### Over Representation Analysis

We perform Over Representation Analysis (ORA), which is based on the fisher’s exact test either on the top/bottom 10% of the ranked (t.value) differential metabolite analysis results or on the clusters from bioRCM. In both cases, we set all detected metabolites as the background.

### Sample metadata Analysis

First, we reduced all the measured features of one sample into a few features in the different principal components (PC) by performing PC analysis (PCA). Since each PC explains a certain percentage of the variance between the different samples, this enables interpretation of sample clustering based on the measured features. By performing anova between each PC and selected patient metainformation (tissue, stage, age, gender), we extracted the main metabolite drivers that separate patients based on these metainformation.

### Visualisation

MetaProViz fine tunes the color palettes, aesthetics, and sizes of elements within each figure layout, with the aim to consistently generate well readable, publication ready figures. For this we use ggplot2-based^26^ figures and adapt figures from EnhancedVolcano^56^, pheatmap^57^ and ComplexUpset^58^.

## DATA AVAILABILITY

Data generated as part of this study are available as supplementary tables or via GitHub: https://github.com/saezlab/MetaProViz/tree/BioRxiv. Information to download all used raw data from the respective studies is available on the “ExampleData” branch of the MetaProViz package (https://github.com/saezlab/MetaProViz/tree/ExampleData).

## CODE AVAILABILITY

The code, release history, reference documentation, and vignettes of the MetaProViz R-package is available at https://saezlab.github.io/MetaProViz/. The code to reproduce all figures and tables in this manuscript is available at https://github.com/saezlab/MetaProViz/tree/BioRxiv.

## ACKNOWLEDGEMENTS

We thank the input and feedback of the members of the Saez-Rodriguez and the Frezza lab, specifically Robin Fallegger, Charlotte Boys, Christoph Mahler, Aurelien Dugourd, Yunfan Bai and Jannik Franken. We also thank all the studies used as example data for making their data publicly available. This work was funded by SmartCare (03LW0233K) to J.S.R. and by the CRUK Programme Foundation award (C51061/A27453) and the Alexander von Humboldt Foundation in the framework of the Alexander von Humboldt Professorship endowed by the Federal Ministry of Education and Research to C.F. D.T. was supported by the Landesinstitut für Bioinformatikinfrastruktur in Baden-Württemberg.

## CONTRIBUTIONS

C.S. developed concepts of MetaProViz R-package and interpreted the biological readouts. D.T. implemented required functionalities within the OmniPath suite. C.S., D.T., D.P. and M.D. developed the MetaProViz R package code. C.S. and D.T. wrote the manuscript. C.S. prepared the figures. All authors reviewed and edited the manuscript. C.F. and J.S.R. supervised the project.

## COMPETING INTERESTS

C.F. is an adviser for Istesso. J.S.R. reports in the last 3 years funding from GSK and Pfizer and fees/honoraria from Travere Therapeutics, Stadapharm, Astex, Pfizer, Grunenthal, Tempus, Moderna and Owkin.

## SUPPLEMENTARY INFORMATION

**Supplementary Figure 1:**
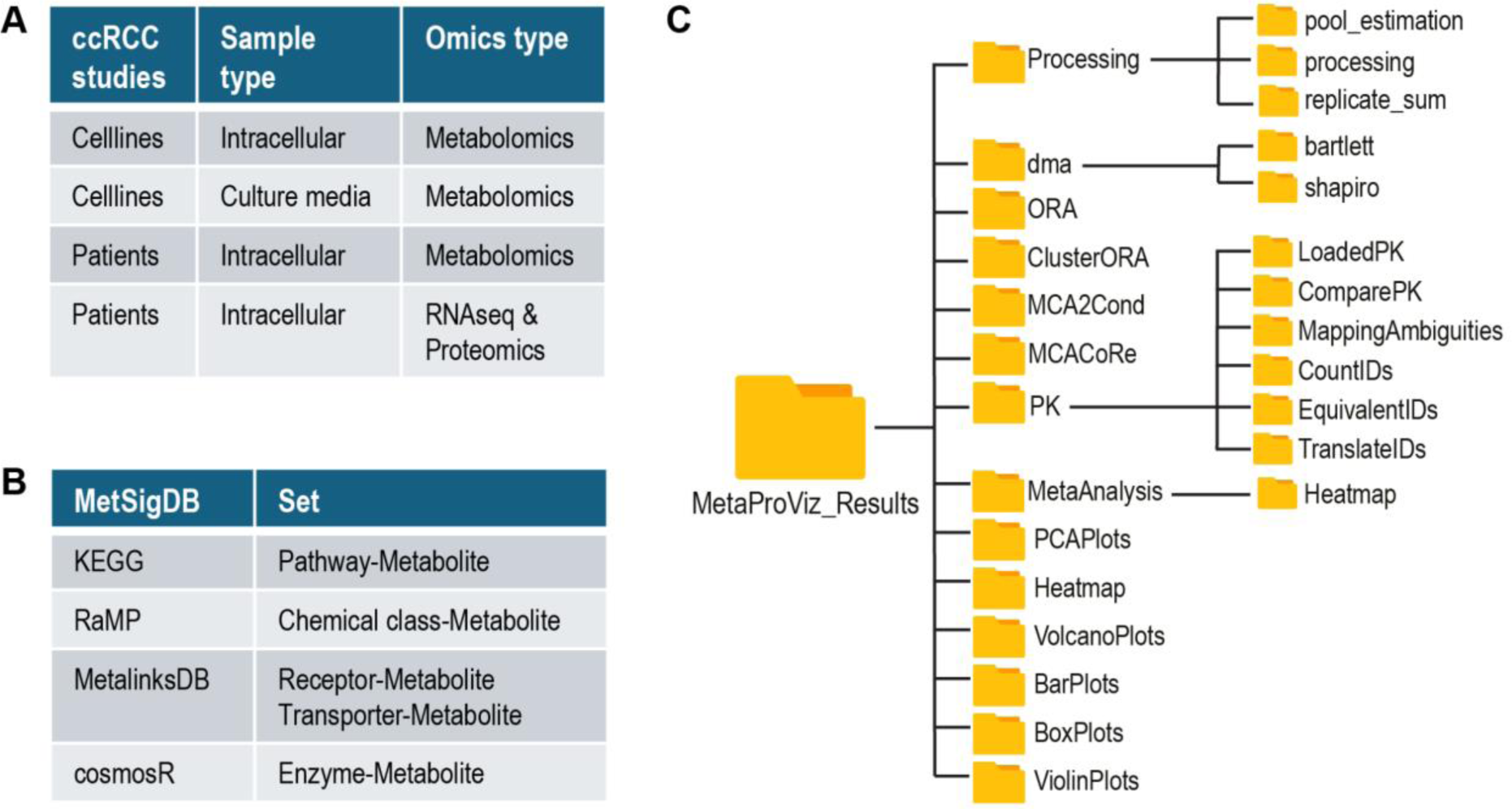
**a)** Overview of available ccRCC example data in MetaProViz. **b)** Output folder structure where the result tables and figures are automatically saved for MetaProViz analysis. **c)** Overview of prior knowledge sets included in MetaProViz MetSigDB collection.

**Supplementary Figure 2:**
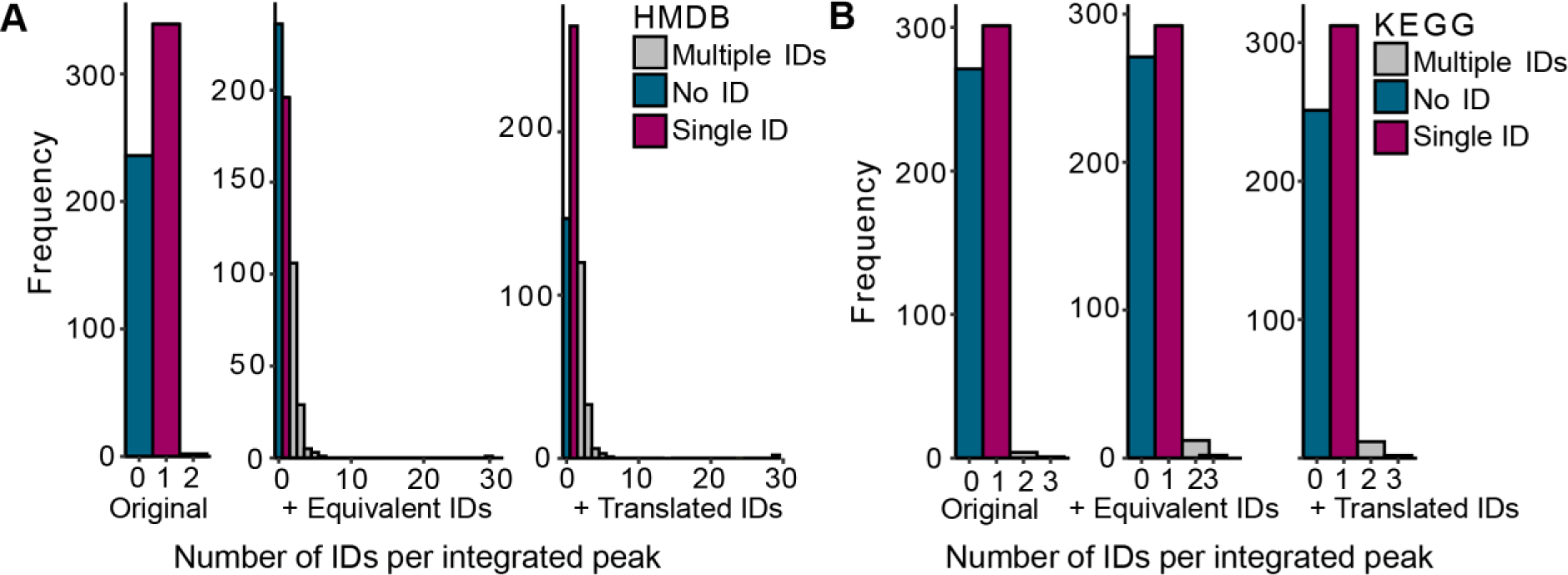
**a-b)** Bargraphs showing the frequency at which a certain number of metabolite IDs per integrated peak are available as per ccRCC patients feature metadata provided in the original publication (left), after potential equivalent IDs where assigned (middle) and after IDs were translated from the other available ID types (right) for **a)** HMDB IDs and **b)** KEGG IDs.

**Supplementary Figure 3:**
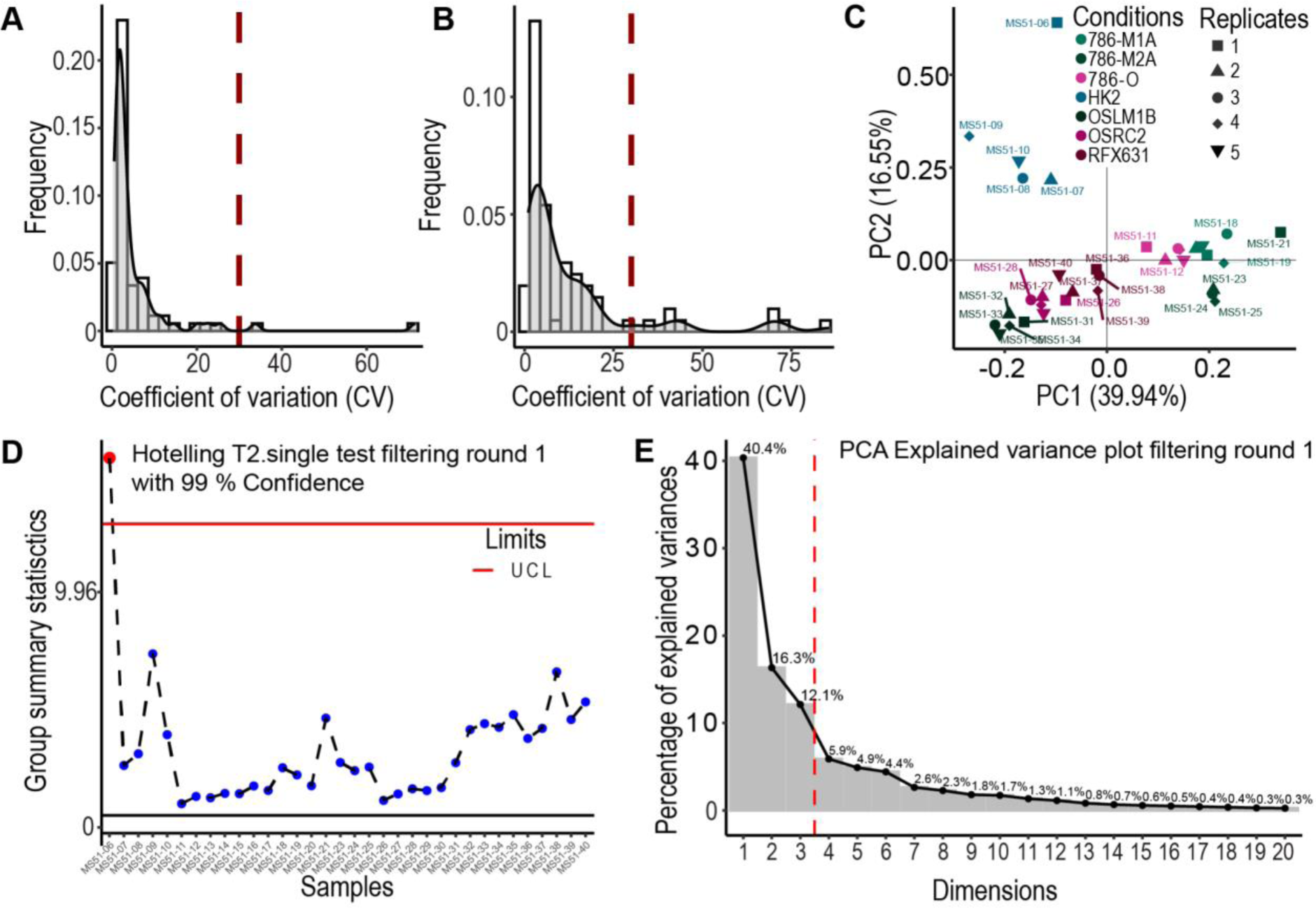
Feature QC and outlier testing of exometabolomics data using MetaProViz. **a-b)** Exometabolomics feature quality of **a)** metabolite pool sample coefficient of variation (CV), **b)** media blank samples, which are samples where no cells were cultured in. **c)** PCA plot with shapes for biological replicates to understand replicate spread. **d-e)** Visualisation of different parameters of Hotelling’s T2 test results with 99% confidence interval of filtering round 1 including **d)** Group summary statistics and **e)** explained variance of PCA plot with elbow (red line). Detailed code, explanation and plots can be found in the vignette (https://saezlab.github.io/MetaProViz/articles/CoRe%20Metabolomics.html).

## Extended data tables

**Extended Data Table 1.**
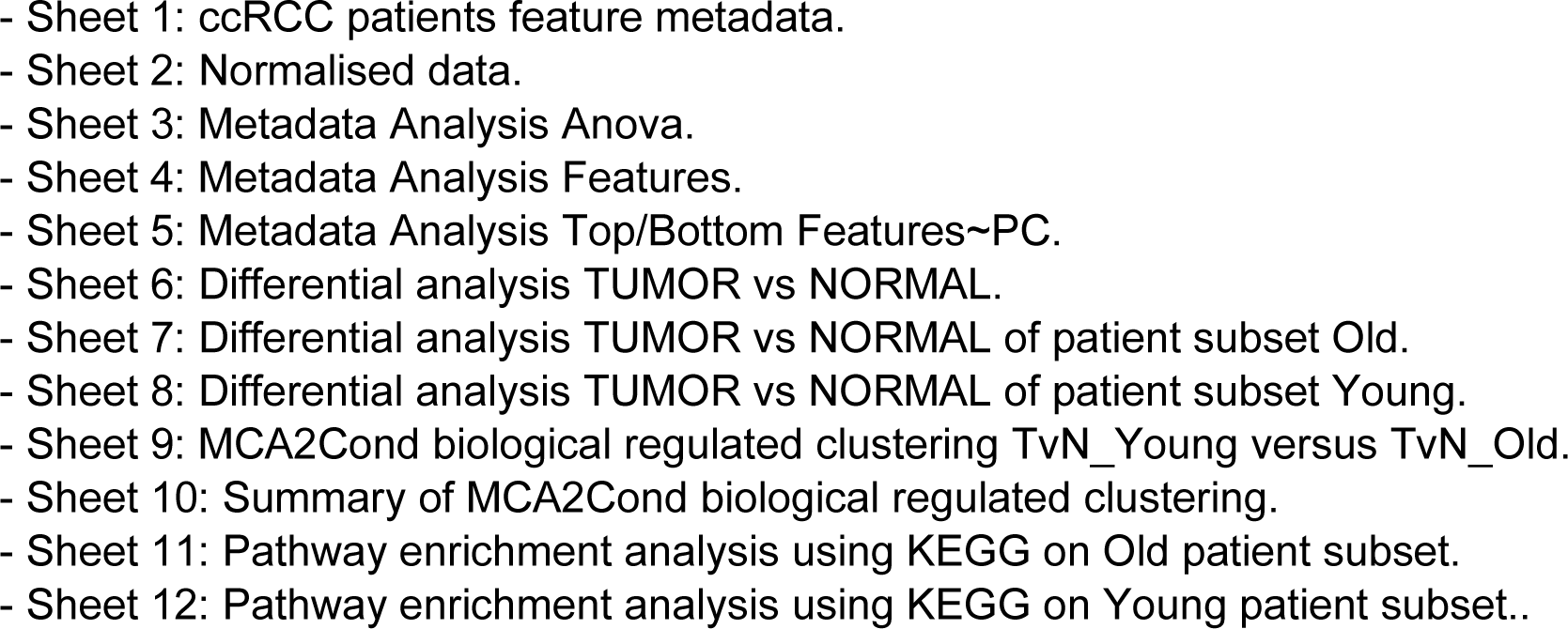
_PatientsTissue_Metabolomics.xlsx:

**Extended Data Table 2.**
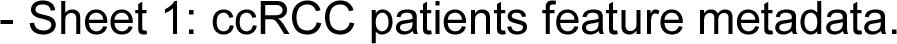
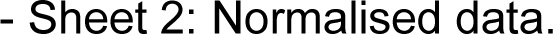
_AlanineMetaboliteIDs.xlsx:

**Extended Data Table 3.**
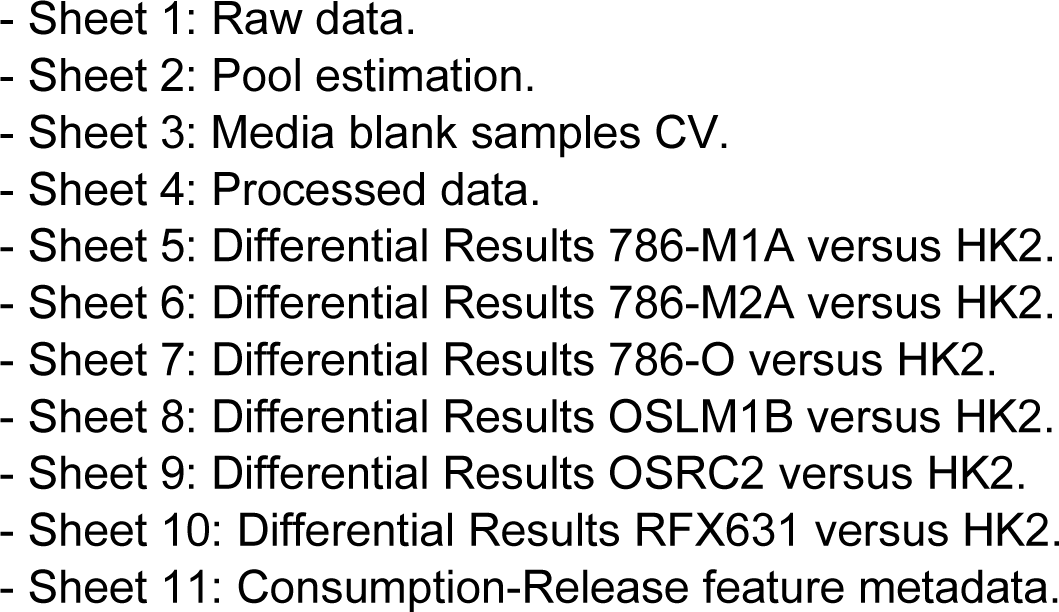
_Cells-Exometabolomics.xlsx:

**Extended Data Table 4.**
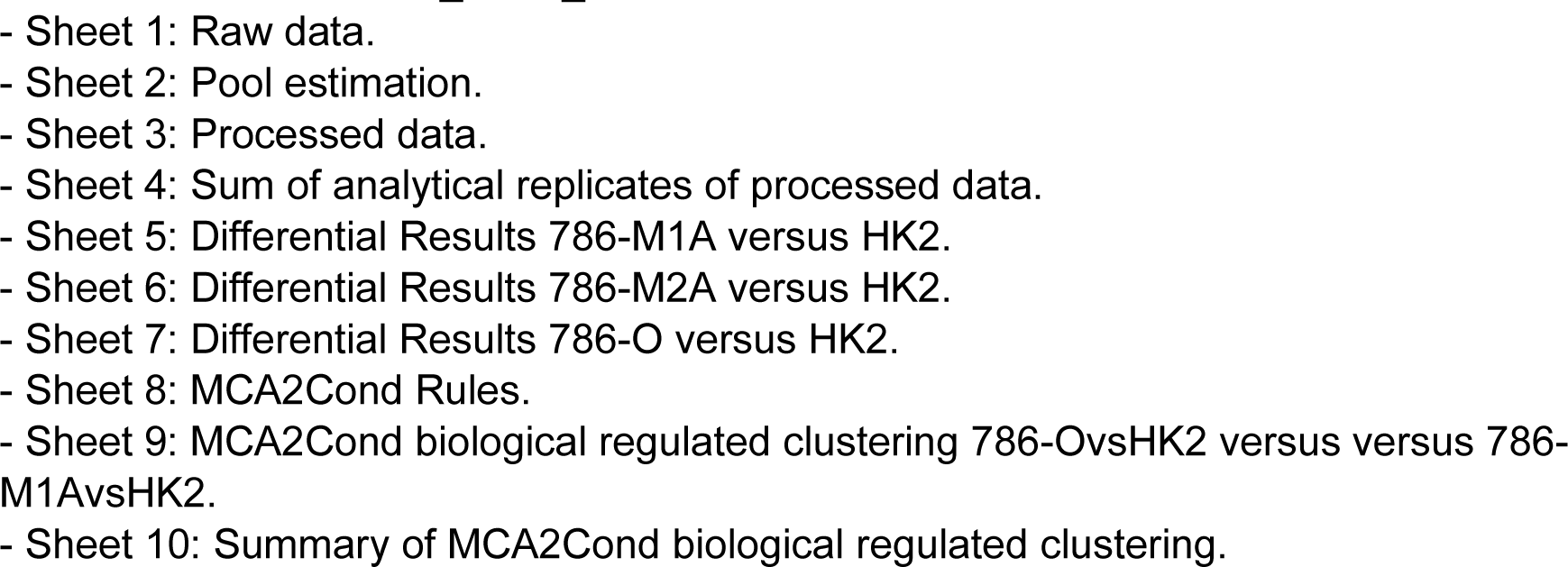
_Cells_Intracellular.xlsx:

**Extended Data Table 5.**
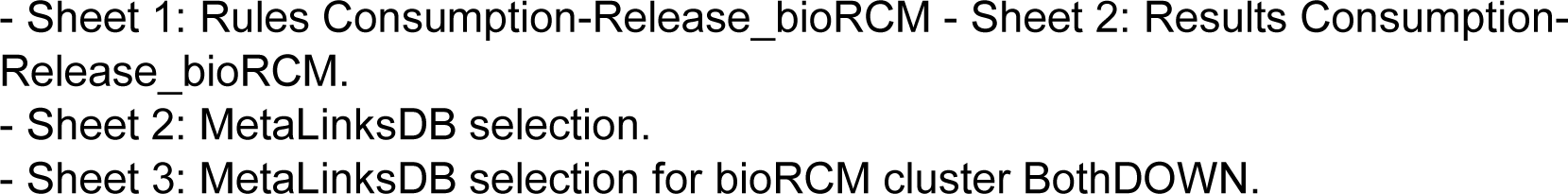
_Integration-extracellular-and-media.xlsx:

## Notes

https://saezlab.github.io/MetaProViz/

https://github.com/saezlab/MetaProViz/tree/BioRxiv

